# Subcortical theta oscillations alternate with high amplitude neocortical population within synchronized states

**DOI:** 10.1101/453837

**Authors:** Erin Munro Krull, Shuzo Sakata, Taro Toyoizumi

## Abstract

Synchronized states are marked by large-amplitude low-frequency oscillations in the cortex. These states can be seen during quiet waking or slow-wave sleep. Within synchronized states, previous studies have noted a plethora of different types of activity, including delta oscillations (0.5-4 Hz) and slow oscillations (<1 Hz) in the cortex and large- and small-irregular activity in the hippocampus. However, it is not still fully characterized how neural populations contribute to the synchronized state. Here we apply independent component analysis (ICA) to parse which populations are involved in different kinds of cortical activity, and find two populations that alternate throughout synchronized states. One population broadly affects cortical deep layers, and is associated with larger amplitude slower cortical activity. The other population exhibits theta-frequency oscillations that are not easily observed in raw field potential recordings. These theta oscillations apparently come from below the cortex, suggesting hippocampal origin, and are associated with smaller amplitude faster cortical activity. Relative involvement of these two alternating populations may indicate different modes of operation within synchronized states.

## Introduction

Cortical state can be described as synchronized or desynchronized (Harris & Thiele, 2011). Desynchronized states are characterized by low amplitude high frequency activity where local neuronal activity is uncorrelated. Desynchronized activity is seen during active waking (engaging in a task such as navigating a maze) and rapid-eye movement sleep (REM). In rats, while the cortex is desynchronized, the hippocampus exhibits theta oscillations, sinusoidal oscillations between 4 and 8 Hz (Diekelmann & Born, 2010). Synchronized states are characterized by high-amplitude, low-frequency oscillations where neuronal populations fluctuate between UP states marked by frequent neuronal firing, and DOWN states marked by neuronal silence. Synchronized states are seen during quiet waking (resting or engaging in a routine task such as eating) as well as non-REM sleep. In rats, while the cortex is synchronized, the hippocampus exhibits irregular activity featuring sharp waves, quickly rising waves accompanied by ripple oscillations (>100 Hz) (Diekelmann & Born, 2010). Within non-REM sleep, cortical activity progresses from K-complexes (large amplitude biphasic waves) to slow-wave activity (<1 Hz) mixed with delta oscillations (1-4 Hz) (V
Crunelli, Cope, & Hughes, 2006). At the same time, the hippocampus can display either low- or high-amplitude irregular activity (Miyawaki, Billeh, & Diba, 2017). While activity during synchronized states have been widely studied, neural ensemble dynamics within the synchronized state is not still fully characterized (Vincenzo
Crunelli, David, Lorincz, & Hughes, 2015; de Andrés, Garzón, & Reinoso-Suárez, 2011a; Diekelmann & Born, 2010; Mizuseki & Miyawaki, 2017; Neske, 2016).

Previous studies on cortical dynamics primarily analyzed cortical activity by looking at either spiking activity or the local field potential (LFP) (Einevoll, Kayser, Logothetis, & Panzeri, 2013). Single- or multi-unit spiking activity gives detailed information about individual neural interactions. However, because the number of neurons we can detect is limited, we must infer how cell populations behave as a whole. On the other hand, LFP recordings reflect activity from many populations simultaneously, including neurons and non-spiking glia (Buzsáki, Anastassiou, & Koch, 2012). However, signals from different populations tend to overlap at each recording site, as illustrated in Fig. 1. The fact that signals overlap makes these recordings difficult to parse, i.e. it isn’t clear which populations are involved in different kinds of cortical activity (Buzsáki et al., 2012; Kajikawa & Schroeder, 2015; Kajikawa & Schroeder, 2011). Current source density (CSD) analysis helps reduce signal spread by focusing on current sinks and sources, but it may still be difficult to parse which cell populations are involved, particularly in the neocortex where neuronal populations are very dense (Gulyás, Freund, & Káli, 2016).

**Figure 1.**
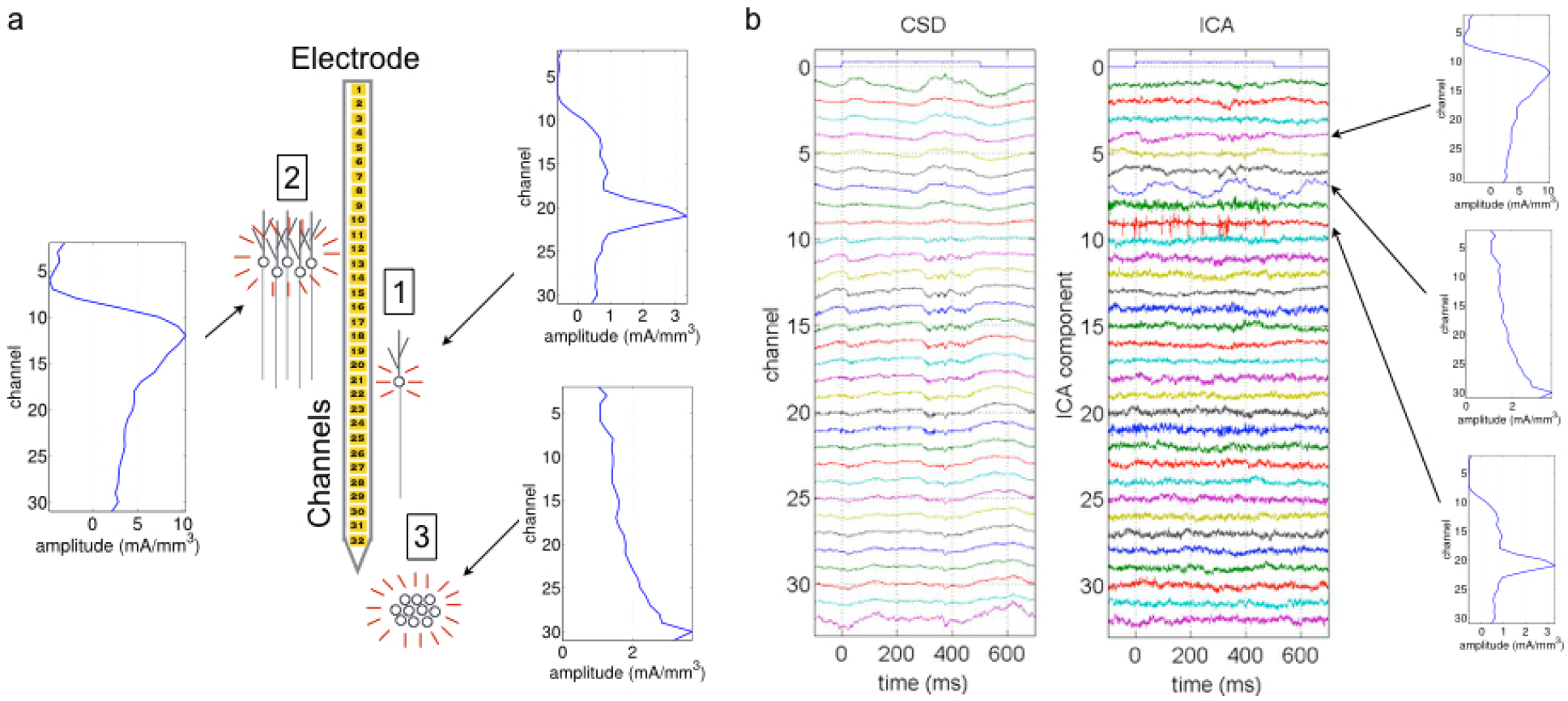
Example of ICA applied to neural data. Data generated from the example anesthetized experiment used for this study. (a) Different populations generate field potentials across the electrode, resulting in overlapping signals recorded at each channel. For example, the signal from a single cell (1) may have the highest amplitude in a single channel, while other channels have much lower amplitude. The signal from a larger population (2) may have high amplitude across multiple channels with a single dipole. A population below the electrode (3) may generate the highest amplitude signal in the channel closest to the population, which doesn’t have a dipole but decays with channels further from the population (Herreras, Makarova, & Makarov, 2015). (b) Example CSD from 32-channel electrode along with ICA decomposition. The ICA components are listed in order from largest to smallest amplitude over the entire experiment. In the CSD, the auxiliary channel 0 indicates a tone is played at time 0, and we see an immediate reaction to the stimulus in channels 11 through 17. This immediate reaction to the tone is summarized in ICA component 4. Each ICA component is associated with a spatial map showing the amplitude of that component across the electrode. The spatial map for ICA component 4 is consistent with the large population (2) in panel a. Similarly, high frequency activity seen in the CSD across channels 11 through 21 is reflected in ICA component 9, whose spatial map is consistent with the single cell (1) in panel a. Finally, there is a 4 Hz sinusoidal oscillation in ICA component 7 that is not apparent in the original CSD. The spatial map gradually increases toward the lowest channel and there is no dipole, indicating that the oscillation was generated from below the electrode as in population (3) in panel a.

To address the issue of overlapping population signals, we turned to Independent Component Analysis (ICA) (Brown, Yamada, & Sejnowski, 2001). Based on the assumption that the LFP is made up of summed population signals, and that the relative amplitude of a single population across the electrode remains fixed over time, ICA decomposes recordings so that signals from different populations are separated into different components. ICA works by finding a matrix that decouples the signals so that they are as statistically independent as possible. Since ICA decouples signals by matrix multiplication, the number of extracted components is the same as the number of recording channels. If there are more populations than recording channels, then ICA groups populations with similar spatial profiles together in the same component. Even with a limited number of channels, ICA has proven to be very effective for parsing signals from different types of neural recordings (Herreras et al., 2015; Jung et al., 2001; Makarov, Makarova, & Herreras, 2010). Moreover, unlike principal component analysis which focuses on high amplitude signals, ICA can easily parse low amplitude signals that may be missed from the raw LFP.

In this study, we applied ICA to neocortical laminar recordings from unanesthetized and urethane anesthetized rats, and consistently observed two ICA components that alternate throughout synchronized states. One component has a high amplitude broad signal that covers much of the lower layers (broad layer 5, BL5); the other component shows intermittent theta oscillations and has a spatial profile indicating it is below the cortex (subcortical, SUB) (Herreras et al., 2015). Furthermore, we found that during synchronized states when BL5 is active cortical activity tends to be slower and have a larger amplitude, while faster small-amplitude activity is seen when SUB is active. These results suggest two modes of operation within the brain during cortical synchronized states.

## Methods

### Animals

We used twenty-three adult Sprague-Dawley rats (male, 270-413 g, n = 18 for anesthetized experiments, n = 5 for unanesthetized experiments). All procedures for anesthetized experiments were performed in accordance with the UK Animals (Scientific Procedures) Act of 1986 Home Office regulations and approved by the Home Office (PPL60/4217 and 70/8883). All procedures for unanesthetized experiments (Sakata & Harris, 2009, 2012) were approved by the Institutional Animal Care and Use Committee of Rutgers University.

### Electrophysiological experiments

Detailed experimental procedures are described in previous studies (Sakata and Harris, 2009, 2012; Sakata, 2016). Briefly, for anesthetized experiments, we anesthetized animals with 1.5-1.6 g/kg urethane. We also administered lidocaine (2%, 0.1-0.3 mg) subcutaneously at the site of incision. After attaching a head-post in the frontal region with bone screws, one of which was used as an electrode for cortical electroencephalograms (EEGs), we placed the animal in a custom head restraint that left the ears free and clear. Two additional bone screws were implanted in the cerebellum as ground. Body temperature was maintained at 37°C with a feedback temperature controller (40-90-8C, FHC). After reflecting the left temporalis muscle, we removed the bone over the left auditory cortex (AC) and carefully performed a small duratomy for each site.

For unanesthetized experiments, in initial surgery, we anesthetized animals with ketamine (100 mg/kg) and xylazine (10 mg/kg), and placed them in a stereotaxic apparatus (David Kopf Instruments). We attached a head-post (Thorlabs, Inc.) with dental cement (3M ESPE, RelyX Luting Cement), removed the left temporal muscle, and covered the exposed bone over the left AC with biocompatible glue and dental cement. After a recovery period, animals were lightly water-deprived, and handling (5-10 min/day) and head-fixation training began. We trained the rats for at least 5 sessions, during which the duration of restraint was gradually extended. Ten percent sucrose was frequently given during training and water was freely available for at least 1 hour after daily training. On the day of recording, we carefully performed a craniotomy and duratomy under isoflurane anesthesia (5% for induction and 0.8% for maintenance). Neither skin nor muscle was cut during this surgery. After a short recovery period (>1 h), recording began.

During recording for both experiments, we covered the brain with 1% agar/0.1 M phosphate buffered saline to keep the cortical surface moist. We inserted a 32 channel silicon probe (A1×32-10mm-50-177-A32, NeuroNexus Technologies) either manually or slowly (2 μm/sec or slower) with a motorized manipulator (DMA-1511, Narishige) into the AC (1400 – 1770 μm from the surface). All electrophysiological experiments were performed in a single-walled soundproof box (MAC-3, IAC Acoustics) with the interior covered by 3 inches of acoustic absorption foam. Broadband signals (0.07–8 kHz) from the silicon probes were amplified (1000 times) (Plexon, HST/32V-G20 and PBX3), digitized at 20 kHz and stored for offline analysis (PXI, National Instruments).

A typical recording schedule was as follows: after insertion of the probe and an additional waiting period (at least 30 min), we started recording with a silent period (at least 5 min), followed by sound presentations, and ended with another silent period (at least 5 min). Acoustic stimuli were generated digitally (sampling rate 97.7 kHz, TDT3, Tucker-Davis Technologies) and delivered in free-field through a calibrated electrostatic loudspeaker (ES1) located approximately 10 cm in front of the animal. We calibrated the tone presentations using a pressure microphone (PS9200KIT-1/4, ACO Pacific Inc) close to the animal’s right ear. Acoustic stimuli for this study consisted of short pure tones (50 ms long with 5 ms cosine ramps, 1/6 or 1/8 octave steps, 3-48 kHz, 10 dB steps, 0-80 dB SPL), long pure tones (300 ms with 10 ms cosine ramps), and unanesthetized experiments also contained brief click trains.

### Histology

After electrophysiological experiments, we perfused rats transcardially with physiological saline followed by 4% paraformaldehyde/0.1 M phosphate buffer, pH 7.4. After overnight postfixation in the same fixative, we incubated brains in 30% sucrose solution for cryoprotection, cut into 100 μm coronal sections with a sliding microtome (SM2010R, Leica), and the sections were collected and placed in 0.1 M phosphate buffered saline (PBS). For verification of silicon probe placement, the free-floating sections were counterstained with NeuroTrace (1/500, N-21480, Life Technologies) in PBS with 0.1% Triton X-100 for 20 min at room temperature. The sections were mounted on gelatin-coated slides and cover-slipped with antifade solutions.

### Data analysis preprocessing

We performed all analysis using Matlab (R2017a, Mathworks, Waltham, MA). Since the sample rate of the original recordings was 20 kHz, we first filtered the data below 650 Hz using the Fast Fourier Transform (FFT, fft in Matlab) with a bump function pass band, where the bump function smoothly transitioned from 0 to 1 over 0.25 Hz. We then removed 50 Hz line noise by linearly interpolating the Fourier transform. Finally, we down-sampled the data to 2 kHz, and high-pass filtered the data over 0.1 Hz using FFT with a bump function pass band.

We noted three kinds of artifacts to remove from analysis: line drift, electrode pops, and epochs with low firing rates. To mark epochs with line drift (large amplitude deviations in the LFP for a single channel), we calculated the standard deviation of each channel. We then marked sections that exceeded 5 standard deviations for that channel. For electrode pops (large jumps in the LFP), we calculated histograms of voltage slopes over the entire experiment for each channel. We marked points where the slope was >2,500 V/s for at least one channel, or >1,000 V/s for at least 2/3 of the channels within 1 ms. To determine overall firing rate, we summed the multi-unit activity (MUA) over all channels, and then computed the average firing rate of the summed MUA over 6 s windows shifted by 600 ms. We marked points that were below 40 Hz for anesthetized experiments and 100 Hz for unanesthetized experiments, where these thresholds were set by the distribution of firing rates for anesthetized and unanesthetized experiments, respectively.

### UP state detection

We used MUA to determine UP and DOWN states. First we calculated the smoothed MUA (sMUA) by summing the MUA over all channels. We then smoothed the summed MUA using a Gaussian filter with standard deviation of 6.25 ms for anesthetized data and 2.5 ms for unanesthetized data (2.5 divided by minimum firing rate). We then computed UP and DOWN state transitions based on Sakata & Harris (2009). The threshold for transitioning from a DOWN to an UP state was the geometric mean of the sMUA over points where sMUA>0, while the threshold for transitioning from an UP to a DOWN state was 1/5 the UP state threshold. The minimum UP/DOWN state length was 50 ms for anesthetized data and 25 ms for unanesthetized data.

### Spectrogram calculation

All spectrograms were calculated over 6 s windows shifted by 600ms using Welch’s power spectral density estimate (pwelch in Matlab). Spectrograms were used to analyze frequency content of the LFP as well as ICA components. To estimate synchronized states, we computed the spectrogram of the LFP for each channel and then took the median over all channel spectrograms, keeping time and frequency fixed. The low-frequency LFP power (LFP-Power) is the summed power from 1-5 Hz (Harris & Thiele, 2011) of the median spectrogram. The threshold for synchronized states for all experiments was set to the median LFP-Power over unanesthetized experiments.

### ICA application

We first spatially filtered the data using a Gaussian kernel with width 50μm. We then calculated the spline inverse CSD over the LFP (Pettersen, Devor, Ulbert, Dale, & Einevoll, 2006). Since the CSD partially separates sources, previous work suggests that ICA separates the CSD more cleanly than the LFP (Lski, Kublik, Swiejkowski, Wróbel, & Wójcik, 2010). For the ICA training data, we randomly selected 100s worth of data points from synchronized data only. We used the extended infomax algorithm to minimize effects of kurtosis in the data (Lee, Girolami, & Sejnowski, 1999) implemented in EEGLAB (Schwartz Center for Computational Neuroscience, La Jolla, CA). We applied ICA 10 times, each on different training data, to yield 10 different sets of ICA components. We then chose the ICA component set that had minimal mutual information between components (Kraskov, Stögbauer, & Grassberger, 2004). To estimate mutual information, we took samples of 100 to 1,000 s of data points. We then calculated a mutual information factor (MIF) by estimating entropy of individual components, using a bins with approximately 10 data points per bin. The MIF is the summed entropy of all individual components, minus log|W| where W is the ICA unmixing matrix.

Out of the 32 components generated by ICA, we selected the SUB component and then the BL5 component. ICA lists components according to amplitude. We chose the first component with a narrow band oscillation between 2-5 Hz in anesthetized experiments, and 5-9 Hz in unanesthetized experiments, reflecting theta ranges in urethane anesthetized and unanesthetized conditions. The BL5 component is the highest amplitude component, not including the SUB component.

ICA component amplitude can vary based on experiment as well as populations included in the component. Therefore, we scaled BL5 and SUB according to their activity levels. We considered BL5 active when its signal was correlated with the sMUA. We calculated the correlation over 6 s windows shifted by 600 ms to match spectrogram data. Since the ICA algorithm guesses the sign of a component based on the spatial profile, we switched the sign of BL5 if necessary so that the correlation was negative overall. Time points with correlation ≥0 were considered inactive. If there were fewer than 30 points that met this criterion, then points with correlation ≥-0.1 were considered inactive. We then centered the signal based on the median and interquartile range of inactive points. To measure the activity level of the SUB component, we first determined the peak frequency between 2-5 Hz for anesthetized data and 5-9 Hz for unanesthetized data. We then smoothed the peak frequencies using a Gaussian with standard deviation 10 s, weighting each point according to the amplitude of the peak. We then found the total amplitude around the peak (width 1 Hz for anesthetized, 2 Hz for unanesthetized), along with the amplitude of the surrounding frequencies which consisted of a half width window above and below the peak window. The activity level of SUB was defined as the scaled peak amplitude, where amplitude was centered on the median and interquartile range of the surrounding frequencies over the entire experiment for anesthetized data. Because surrounding frequency amplitude varied a great deal over the course of unanesthetized experiments, we decided to scale the peak amplitude based and median and interquartile range over 6s instead of the whole experiment.

### Sleep stage estimation

We estimated sleep stages only on the LFP median spectrogram for unanesthetized experiments. We first calculated the LFP delta (0.5-4 Hz), theta (6-9 Hz), and sigma (10-14 Hz) power (Benington, Kodali, & Heller, 1994; Gross et al., 2009). We then smoothed the power using a Gaussian kernel with width 2.5 s. As sigma*theta power indexes waking vs. sleeping, we selected the sigma*theta threshold first based on largest population density (Benington et al., 1994). We calculated a histogram with 100 bins over the lowest 90% of data points. Then we smoothed the histogram using a Gaussian with width over 1 bin. We calculated the max height of the histogram h_c_ and mean height of the histogram h_m_. The threshold is the first point above h_c_ where the histogram falls below (h_c_+h_m_)/2 as sigma*theta increases. The delta/theta threshold, which signifies the border between REM and non-REM sleep, was based on the non-REM to REM indicator value (NIV) (Benington et al., 1994). We calculated candidate transition to REM (NRT) segments where delta power dropped (delta-DP<1 in (Benington et al., 1994)) and sigma*theta is above threshold. For each NRT segment, we calculated the NIV and the minimum delta/theta value over the entire segment (DT-min). The threshold was set to the median DT-min over all segments with NIV values in the top 50%.

### Laminar estimation

To display channel data in Figure 6a, we estimated the central L4 channel based on the maximum CSD amplitude evoked by the preferred tones. The channel with the highest amplitude was labelled as a thalamic recipient layer (L4).

### Statistical analysis for cortical activity

To compare cortical activity across states, we used 2-way ANOVAs based on conglomerated artifact-free data points across all anesthetized or unanesthetized experiments with BL5/SUB index *φ* between −45 and 135 degrees. All 2-way ANOVAs were performed on the variable of interest as a function of LFP-Power and *φ*, where LFP-Power was considered continuous and *φ* was split into BL5 (*φ* ≤45) and SUB (*φ* >45). For UP state location, we also included the experiment as a random factor.

UP phase amplitude was defined as the largest amplitude occurrence after Gaussian smoothing over time with a kernel of 2ms. Peak frequency was defined as the maximum frequency after Gaussian smoothing with a kernel of 0.25 Hz. Peak frequency width was determined by the Matlab function findpeaks.

### Data and code availability

All data analyzed in this study is available upon request from the corresponding author. Any code used to analyze the data is also available upon request.

## Results

We used ICA to investigate which cell populations are active during synchronized states in 18 urethane anesthetized and 5 unanesthetized head-fixed rats. Anesthetized rats exhibited sleep-like states (Clement et al., 2008; Pagliardini, Gosgnach, & Dickson, 2013), while unanesthetized rats could be waking or sleeping. We recorded from all layers of the auditory cortex using 32-channel linear probes (Sakata & Harris, 2009, 2012). From these recordings, we calculated the CSD before applying ICA in order to get clearer components (Lski et al., 2010). We applied ICA to the CSD 10 times, each time with different randomly selected sets of artifact-free training data, and chose the ICA run with the least estimated mutual information between components (Kraskov et al., 2004) to promote as much separation as possible.

### Two populations consistently revealed by ICA

We consistently noticed two ICA components. One component exhibited sinusoidal oscillations in the theta range (Buzsáki, 2002; Buzsáki & Draguhn, 2004; Lakatos et al., 2005; Montgomery, Betancur, & Buzsaki, 2009) that appeared to be subcortical (SUB). The other was the highest amplitude component which broadly affected all layers, centered on layer 5 (BL5).

We define the SUB component as the highest-amplitude component with clock-like sinusoidal oscillations in the theta range (2-5 Hz anesthetized, 5-9 Hz unanesthetized). We found a SUB component in 16/18 anesthetized experiments and 5/5 unanesthetized experiments. Figure 2a shows that the amplitude of the SUB component gradually increased toward the lowest channel in anesthetized experiments, suggesting that signals originated from below the cortex (Herreras et al., 2015). In unanesthetized experiments, the amplitude was even over all channels. At the same time, we noticed that signals associated with motion, which affects all channels evenly, were also separated into the SUB component. Note that motion-associated signals may be present outside time intervals marked as motion artifact (see Methods). Therefore, it is possible that ICA grouped SUB and motion signals together because of their similar spatial maps.

**Figure 2.**
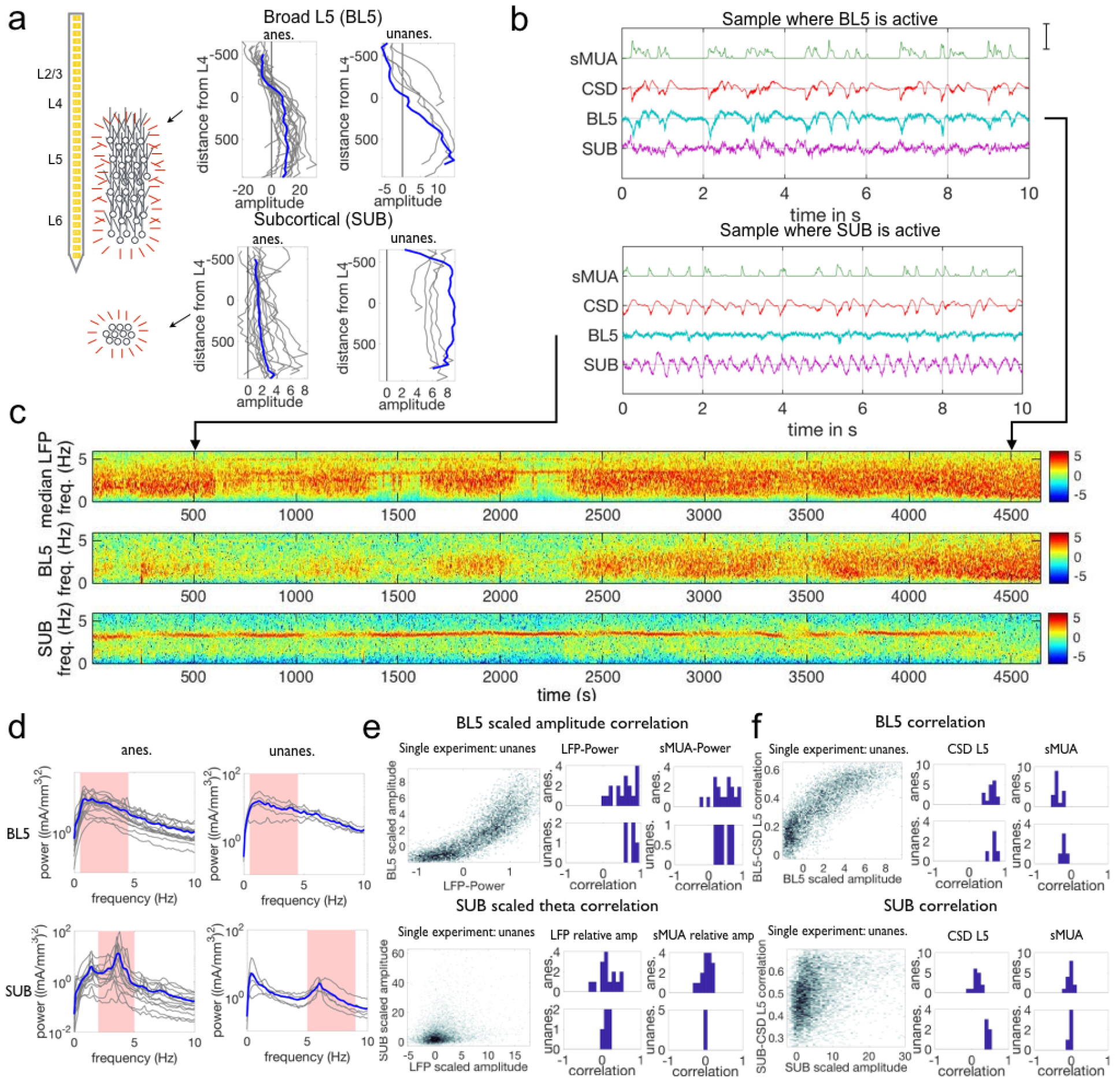
Properties of BL5 and SUB components. (a) Spatial maps of both components across 32 channels used to record the LFP, along with diagram of putative population sources. Results for the example anesthetized experiment shown in (c) and unanesthetized experiment in Supplementary Fig. 1 are highlighted in blue throughout this study. Spatial maps indicate BL5 is made up of a larger population located within the cortex, while SUB is generated by a population below the cortex. Amplitude in mA/mm^3^, distance in μm. (b) Segments when BL5 is active (strong low-frequency oscillations) and when SUB is active (strong sinusoidal oscillation of approximately 4 Hz). Note that the CSD and smoothed multi-unit spiking activity (sMUA) show low-frequency oscillations in both cases. Scale bar is 85th percentile amplitude of sMUA, CSD in channel 17, and ICA component taken from channel with maximum amplitude in spatial map. (c) Spectrograms of the LFP, BL5 and SUB components for an example anesthetized experiment. (d) Average frequency spectra when BL5 and SUB are active (scaled amplitude ≥2). Frequencies used to determine activity level are highlighted in pink. (e) Correlation between BL5 scaled amplitude with LFP-Power and sMUA low frequency power (1-5 Hz), and SUB scaled amplitude with LFP and sMUA relative theta amplitude. Density plot for example experiments on the left, histograms of correlation from density data on the right. (f) Correlation between BL5/SUB signal and CSD from L5 (channel 350 μm below L4) and sMUA. Example experiments on left, histograms of median correlations over points where BL5/SUB are active on the right.

We define the BL5 component as the largest amplitude component which was not the SUB component. Because BL5 was simply the largest component, each experiment had a BL5 component. Spatial maps of BL5 consistently had large amplitude across all lower layers, with the largest amplitude in L5 and a dipole in L4. These spatial maps are consistent with a large population generating the signal across multiple channels as illustrated in Fig. 2a.

BL5 and SUB were active intermittently, as shown in the example anesthetized experiments in Fig. 2b. When BL5 was active, it exhibited large amplitude low frequency oscillations that closely resembled the CSD. Because of this, we define the BL5 amplitude based on the amplitude from 0.5-4.5 Hz (anesthetized) and 0.5-7 Hz (unanesthetized). When SUB was active, it exhibited a sharp peak frequency between 2-5 Hz (anesthetized) or 5-9 Hz (unanesthetized). Fig. 2c shows the average frequency spectra of both BL5 and SUB when they are active, as defined below.

In order to compare activity level across experiments, we scaled the BL5 and SUB amplitude according to the noise level inherent in their signals. We found that BL5 has a high correlation with the CSD and smoothed Multi-Unit Activity (sMUA) when active, and was uncorrelated with CSD and sMUA otherwise. Therefore, we scaled the BL5 component according to its amplitude when correlation with sMUA is low (see Methods). Likewise, in the SUB spectrogram in anesthetized experiments, there was a sharp theta peak which stood out among other frequencies when SUB was active, and blended in with surrounding frequencies when it was not. We therefore scaled SUB amplitude according to the amplitude of the surrounding frequencies over the entire experiment (see Methods). In unanesthetized experiments, while the theta peak stood out during desynchronized states and intermittently during synchronized states, the surrounding frequencies changed according to overall low-frequency LFP power (LFP-Power, 1-5 Hz power of median spectrogram over all channels). (See Supplementary Fig. 1.) In order to not overestimate the SUB amplitude during synchronized states, we scaled SUB amplitude according to surrounding frequencies within a 6 s window. Since both BL5 and SUB are scaled according to the median and interquartile range of their inherent noise level, we consider BL5 and SUB to be active if their amplitude is ≥2.

Since the spatial maps indicate that BL5 has cortical origin while SUB originates from below the cortex, we investigated the relationship between BL5 and SUB and the local activity. First, we found the correlation between the BL5 scaled amplitude and LFP-Power as well as sMUA 1-5 Hz amplitude (sMUA-LFA) (Fig. 2d). We found that the BL5 scaled amplitude had a positive correlation with both LFP-Power (t-test, anesthetized: n=17, p<0.0001, unanesthetized: n=5, p<0.0001) and sMUA-LFA (t-test, anesthetized: n=17, p<0.0001, unanesthetized: n=5, p<0.0062), indicating that BL5 is linked with local spiking activity. (See Supplementary Table 1 for details on statistical tests.) Using the scaled theta amplitude of the LFP and sMUA signals, computed in the same way as the SUB component, we calculated the correlation between the SUB scaled amplitude and the LFP and sMUA scaled theta amplitude. SUB scaled amplitude was not correlated with sMUA scaled theta amplitude (t-test, anesthetized: n=15, p=0.4623, unanesthetized: n=5, 0.5536) giving further evidence that SUB is not cortical. At the same time, SUB scaled amplitude showed some correlation with LFP scaled theta amplitude (t-test, anesthetized: n=15, p=0.0260, unanesthetized: n=5, p=0.0532) indicating that theta oscillations were detectable in the LFP.

While the above results confirm that the BL5 and SUB scaled amplitude align with the hypothesis that BL5 has cortical origin while SUB originates below the cortex, our measure of BL5 and SUB activity levels only take the overriding frequency of their signal into account. Therefore, we decided to investigate in more depth by directly taking the correlation of the BL5 and SUB signals with the CSD from L5 and sMUA (using 6 s windows shifted by 600 ms in the same manner as the spectrogram). We found that the median correlation between BL5 and the CSD or sMUA when BL5 is active was significant overall (t-test, anesthetized: n=17, p<0.0001, unanesthetized: n=5, p<0.0001 for CSD and anesthetized: n=17, p<0.0001, unanesthetized: n=5, p=0.0027 for sMUA), further attesting involvement with local cortical activity. The correlation between SUB and sMUA was not significantly far from zero (t-test, anesthetized: n=15, p=0.4033, unanesthetized: n=5, p=0.1243), confirming that SUB had little relation with local spiking activity. At the same time, the correlation between SUB and CSD was positive (t-test, anesthetized: n=17, p<0.0001, unanesthetized: n=5, p<0.0001), showing involvement with SUB in the CSD. Overall, while BL5 showed strong correlation with local activity, which is consistent with the interpretation that the BL5 population is cortical. SUB showed little correlation with local spiking activity, confirming the SUB population is not local. At the same time, SUB was correlated with the LFP and CSD, indicating it has significant influence on local field recordings.

### SUB amplitude alternates with BL5 amplitude

Fig. 2b appears to show that SUB and BL5 alternate with each other. To test this possibility, we calculated the correlation between the BL5 and SUB scaled amplitudes as illustrated in Fig. 3ai, and found that they were anticorrelated over all experiments (Fig. 3aii, t-test, anesthetized: n=15, p=0.0002, unanesthetized: n=5, p=0.0054). (See Supplementary Table 2 for details on statistical tests.) Because BL5 and SUB are anti-correlated, we quantified the relative BL5-SUB participation in the CSD by calculating the angle *φ* and radius r for each BL5-SUB data point as shown in Fig. 3ai. In the conglomerate density plots of *φ* p vs. r over all experiments (Fig. 3aiii), we can see peak densities where BL5 is dominant (*φ* ≤45) and where SUB is dominant (*φ* >45). Thus, it makes sense to refer to BL5- and SUB-dominant states.

**Figure 3.**
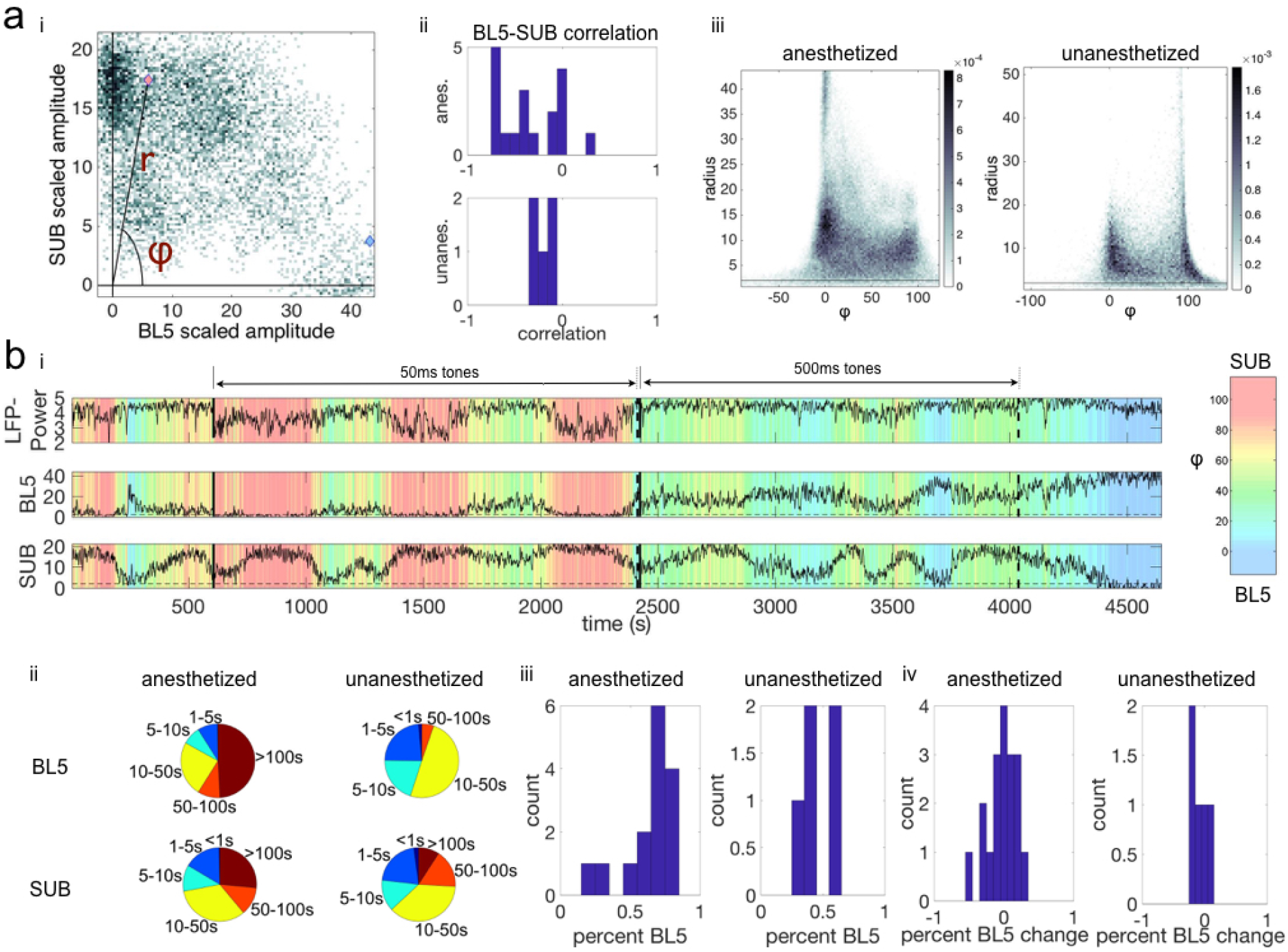
BL5 and SUB alternate within experiments. (a) i. Density plot of BL5 vs. SUB scaled amplitude for example anesthetized experiment (same as Fig. 1b) Pink and blue marks indicate points that are used for example BL5 and SUB active segments in Fig. 1b, respectively. We set φ as the angle from the BL5-axis, while r is the radius. ii. Correlations of BL5 vs. SUB scaled amplitude for each experiment. iii. Conglomerate density plots of φ vs. radius for all anesthetized and unanesthetized experiments. (b) i. Time course of φ over the same example anesthetized experiment used in Fig. 1. ii. Pie charts showing the amount of time spent in BL5 vs. SUB-dominant episodes. For example, for anesthetized experiments about 1/3 of time spent in SUB is in an episode of length 5-10s. iii. Histogram of the percentage of time spent in BL5-vs. SUB-dominant states. iv. Histograms of the difference in percentage of time dedicated to BL5-dominant states during tone presentations vs. spontaneous activity.

We then looked into how long BL5- and SUB-dominant states last. Fig. 3bi illustrates how *φ* changes over the entire example anesthetized experiment, where global trends tend to switch on the order of 100s of seconds with finer-grained changes occurring throughout the experiment. In Fig. 3bii, we see that although BL5-dominant states tend to last longer, both states can last on the order of 10-50s. The amount of time spent in a BL5-dominant vs. SUB-dominant state was similar (Fig. 3biii, mean and standard deviation 0.6283±0.1656 for anes, 0.4659±0.0871 for unanes.). Furthermore, tones played during experiments do not appear to affect the length of states (Fig. 3biv, percent tone vs. spontaneous activity 0.0165±0.1839 for anes., −0.0791±0.1472 for unanes.).

### BL5 and SUB amplitudes alternate during synchronized states

In Fig. 2b, we see instances where either BL5 or SUB are active in the presence of noticeable CSD oscillations as seen during the synchronized states. Synchronized states were previously characterized by high LFP-Power (Harris & Thiele, 2011). Since we showed in the above section that BL5 and SUB alternate within experiments, we further investigated whether BL5 and SUB alternate within synchronized states as well. Fig. 4a shows the density plot of LFP-Power vs. *φ* for the same example experiment as Fig 2b. Within this plot, *φ* ranges from 0 to 80 degrees for higher values of LFP-Power in this experiment. Calculating the range (5-95 percentile) over the density for all experiments reveals a wide range of *φ* where both BL5- and SUB-dominant states are included in the same echelons of LFP-Power. In contrast, *φ* often took high values (>45) when the LFP-Power is low.

**Figure 4.**
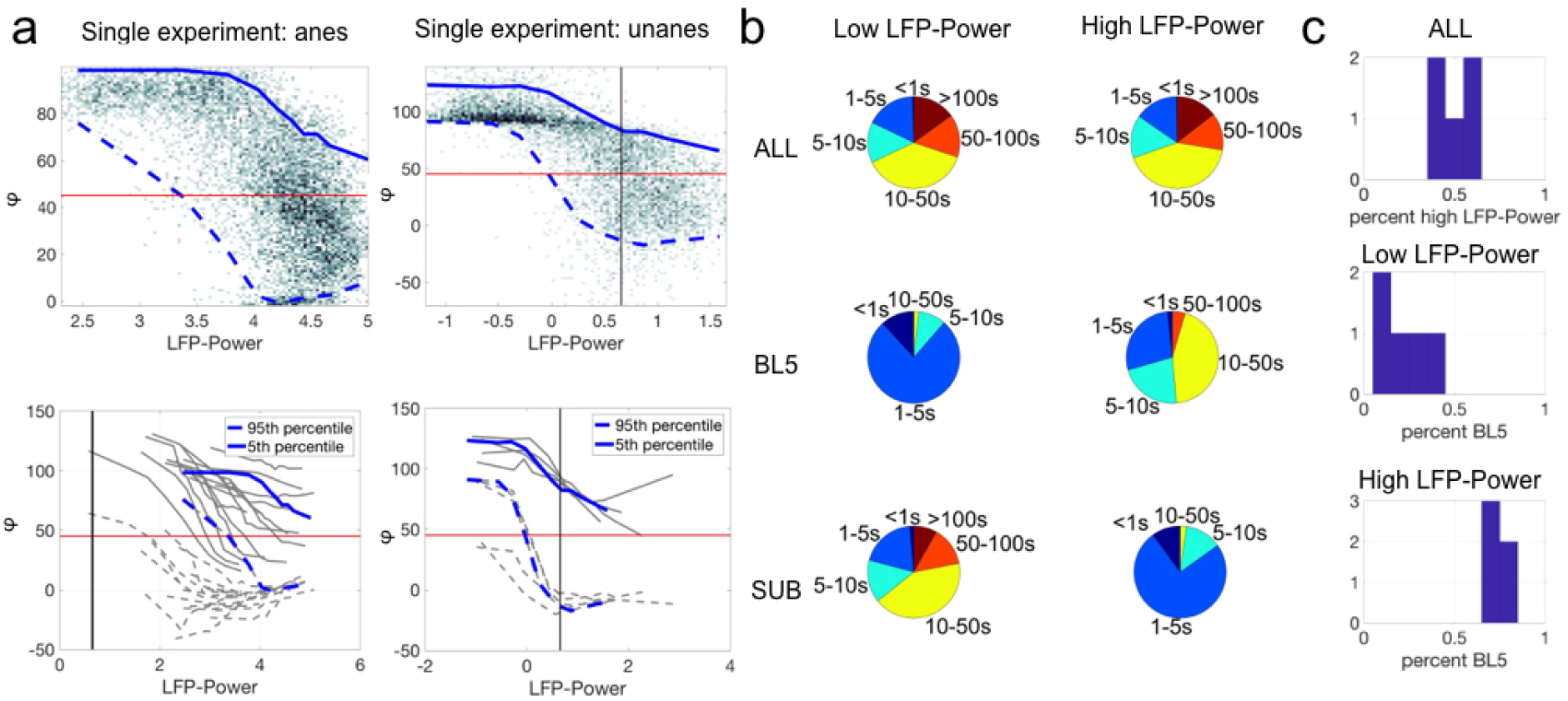
BL5 and SUB alternate within synchronized states. (a) Top plots show density of LFP-Power vs. φ for single experiments along with 5-95 percentile ranges. Bottom plots show ranges for all experiments with results for example experiments highlighted in blue. Vertical black lines indicate median LFP-Power over unanesthetized experiments, used as the threshold between low vs. high LFP-Power. (b) Distribution of episode lengths as a percentage of time spent in episodes of that length. (c) Percentage of time spent in each state. The top panel shows the percentage of time spent with high LFP-Power vs. low LFP-Power. The lower two panels show the percentage of time spent in BL5 vs. SUB-dominant states. Notably, approximately 30% of time in high LFP-Power is SUB-dominant. Unanesthetized experiments only.

To see how much time is spent in BL5-vs. SUB-dominant states within episodes with low vs. high LFP-Power (threshold defined as the median over all unanesthetized experiments), we calculated the duration of each episode along with the percentage of time spent in BL5-vs. SUB-dominant states. We found that while BL5-dominant states were considerably longer than SUB-dominant states when LFP-Power was high, the total amount of time spent in BL5- and SUB-dominant states were both significant, with about 70% of time spent with BL5-dominant vs. 30% with SUB-dominant, similar to the ratio found in Fig.3biii (Fig. 4bc, mean ± standard deviation: 0.5033±0.0949 percent high LPF-power, 0.1906±0.1158 percent BL5 during low LFP-power, 0.7373±0.0334 percent BL5 during high LFP-power). In contrast, the majority of the time was spent in SUB-dominant states when LFP-Power was low. We further investigated the relationship between SUB theta oscillations and LFP-Power directly, since theta oscillations were previously noticed predominantly during desynchronized states, and not synchronized states (Diekelmann & Born, 2010). Supplementary Fig. 2 reveals a similar relationship between LFP-Power and SUB theta oscillations, with a wide range of amplitudes over synchronized states. Furthermore, the length of episodes with significant SUB scaled amplitude during synchronized states is approximately the same as episodes without significant SUB activity. Likewise, about 50% of the time spent in synchronized states had significant SUB scaled amplitude (Supp. Fig. 2c, mean ± standard deviation: 0.3716±0.2059 percent low SUB for anes. 0.3486±0.0997 for unanes., 0.2331±0.1166 percent low SUB during low LFP-Power and 0.4698±0.0653 percent low SUB during high LFP-Power for unanes.).

### BL5 and SUB amplitudes alternate within sleep parameters

Synchronized states occur during both waking and sleeping. To further characterize the occurrence of BL5-vs. SUB-dominated states, we compared their amplitudes with traditional measures of sleep state. In particular, we calculated the delta (0.5-4 Hz), theta (6-9 Hz), and sigma (10-14 Hz) power for each unanesthetized experiment. We then compared *φ* p with the measures sigma*theta, used to indicate wake vs. sleep, and delta/theta, used to indicate non-REM vs. REM sleep (Benington et al., 1994; Gross et al., 2009). Fig. 5a shows how *φ* changes for a single unanesthetized experiment with respect to LFP-Power, delta/theta, and sigma*theta. Although the first few episodes with high LFP-Power and sigma*theta appear to be BL5-dominated, the last episode appears to have many points where SUB is dominant. We then compared BL5-dominated vs. SUB-dominated points directly on the delta/theta and sigma*theta plane. In Fig. 5b, we can see that the top right quadrant which indicates non-REM sleep has clear overlap of both BL5 and SUB points. As expected, we predominantly see SUB-dominant states when delta/theta or sigma*theta is low. Furthermore, a summary of state durations along with percentage of time spent in each state indicate that while episode with BL5-dominant states tend to be longer within the estimated non-REM sleep region, a significant amount of time is spent in both BL5- and SUB-dominant states (Supp. Fig. 3, mean ± standard deviation percent BL5: 0.0505±0.0535 during active wake, 0.0976±0.0885 during quiet wake, 0.3062±0.1447 during REM, 0.6846±0.0540 during non-REM). Remarkably, approximately 70% of time is spent with BL5 dominant while 30% of time with SUB dominant, similar to high LFP-Power states. At the same time, approximately 50% of time within the estimated non-REM region has significant SUB theta amplitude (Supp. Fig. 3, mean ± standard deviation percent low SUB: 0.1210±0.1030 during active wake, 0.1832±0.1551 during quiet wake, 0.2623±0.0734 during REM, 0.4651±0.0679 during non-REM).

**Figure 5.**
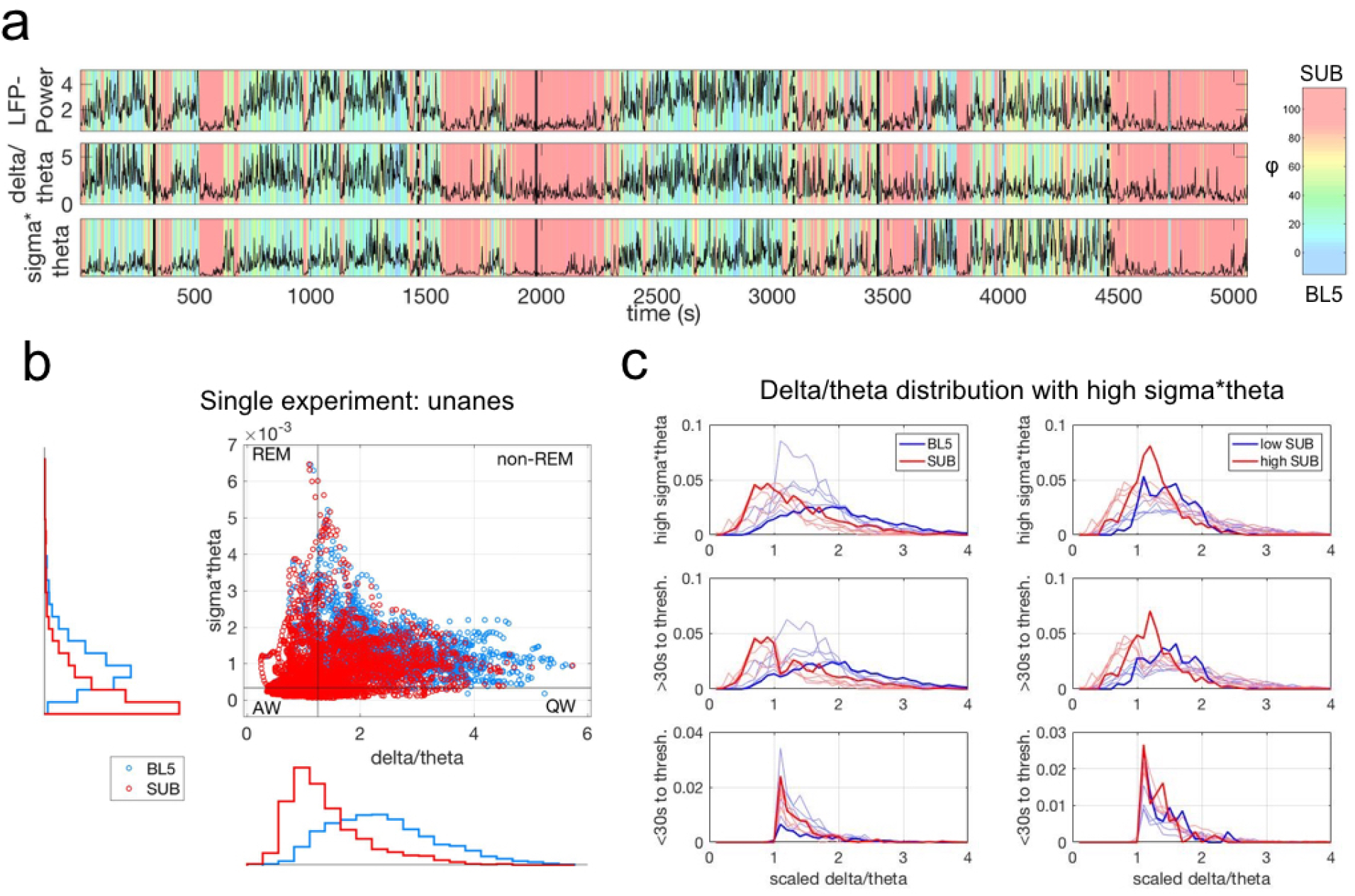
BL5 and SUB dominant states alternate within traditional sleep parameters. (a) Phi transitions with respect to LFP-Power, delta/theta power, and sigma*theta power for the same unanesthetized experiment as Supp. Fig. 1. (b) SUB-dominant points (φ >45) mapped on top of BL5-dominant points (φ≤45) on the delta/theta vs. sigma*theta plane used to help distinguish sleep stages. The upper part of the plane indicates sleep, while the lower part indicates waking. High delta/theta indicates synchronized states such as non-REM sleep, while low delta/theta indicates desynchronized states such as REM sleep. Black boundaries indicate thresholds used for panel c. (c) Histograms of BL5 vs. SUB points on the left, and high SUB vs. low SUB amplitude on the right. Delta/theta is scaled by the delta/theta threshold for each experiment. Upper plots include all points above the sigma*theta threshold, set to be just above the peak sigma*theta density. Middle plots include points at least 30s away from a transition across the delta/theta threshold. Lower plots include points within 30s of a delta/theta threshold crossing. Results for example experiment highlighted in bold.

One reason for seeing theta during non-REM sleep could be the transition to REM state, where theta oscillations become strong just before REM (Benington et al., 1994; Gottesmann, 1996). To investigate whether SUB theta could represent the transition to REM, we plotted histograms of BL5 and SUB points for sigma*theta above threshold (see Methods), shown in Fig. 5c. We then removed points in the 30s leading up to the delta/theta threshold signifying a transition to REM. The histograms show that BL5- and SUB-dominated points have a similar distribution within the estimated non-REM region of the sigma*theta vs. delta theta plane when transition points are removed. We furthermore checked the relationship of SUB theta amplitude with these sleep parameters directly. When we removed transitional points from the non-REM region, we also found similar distributions between high SUB theta amplitude vs. low SUB theta amplitude points. Together, these figures suggest that SUB-dominant states do not simply comprise transition-to-REM states. versions of slow oscillatory activity in the L5 CSD and sMUA, one where BL5 is dominant and one where SUB is dominant. Looking at the CSD and spiking activity across cortical layers in Fig. 6a, we also notice qualitative differences in the activity between BL5 and SUB-dominant states. In particular, we noticed that BL5 states tended to have larger UP states centered in L5 with a slower frequency. At the same time, SUB states tended to have smaller UP states focused in L4 with a higher frequency.

**Figure 6.**
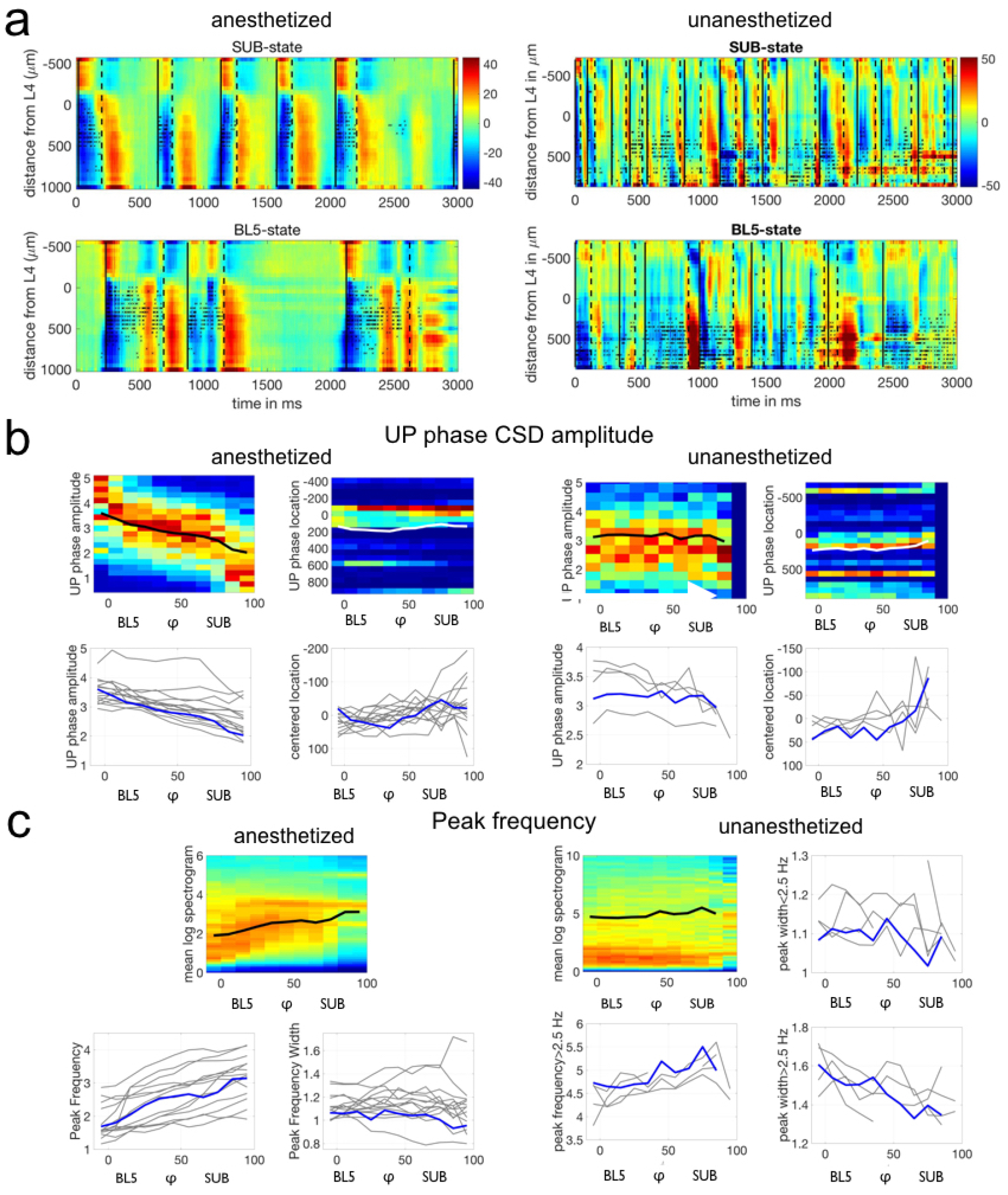
Neocortical activity differs depending on BL5-vs. SUB-dominant states. (a) CSD (color, mA/mm^^^3) and spiking activity (black asterisks) from example anesthetized and unanesthetized experiments seen in previous figures. UP state onsets are marked by solid black lines, while DOWN state onsets are marked by dashed lines. (b) Top plot shows density of UP state amplitude and UP state amplitude peak location for individual experiments, along with the mean with respect to φ. Density is normalized for each φ bin, and mean for a φ bin is calculated only if there are ≥30 points. We only included synchronized data points. Lower plots show the mean for all experiments, with highlighted results for the example experiment above. (c) Top left plot shows average spectrogram values over φ for the same example experiments as above, along with the mean peak frequency over φ. Other plots show means for all experiments, with highlighted results for the example experiment.

To see if UP state CSD amplitude systematically changed with *φ*, we compiled the amplitude of all UP states during synchronized states (LFP-Power greater than median LFP-Power over all unanesthetized experiments). We noticed a down-ward trend in amplitude across *φ* in each experiment (see Fig. 6b). We then tested whether UP state amplitude decreased with respect to *φ* independently from LFP-Power, because LFP-Power may also predict amplitude changes in individual UP states. We found that UP state CSD amplitude significantly decreased with *φ* beyond LFP-Power (2-way ANOVA with LFP-Power and *φ* conglomerated over experiments, anesthetized: p<0.0001, unanesthetized: p<0.0001). (See Supplementary Table 3 for details on statistical tests.) Since UP states appeared to be stronger in L5 during BL5-dominant states, we also tested the UP state amplitude peak cortical depth. Similarly, we found that UP state cortical depth decreased with respect to *φ* beyond LFP-Power (2-way ANOVA between LFP-Power and *φ* with experiment as random variate, anesthetized: p<0.0001, unanesthetized: p=0.0023).

Finally, we investigated how the frequency content of the LFP changes with respect to *φ*. Figure 6c shows that the peak frequency tends to increase for anesthetized experiments, and the peak in the spectrogram narrows indicating that oscillations become more regular. We found that this trend is significant over all experiments independent of LFP-Power (2-way ANOVA between LFP-Power and *φ*, p=0 for peak frequency, p=0.008 for peak frequency width).

The frequency content of unanesthetized experiments proved to be richer than anesthetized experiments, with strong low frequency oscillations throughout synchronized states and an increasing envelope of delta frequencies. We tested whether the envelop of delta frequencies increased for all experiments by measuring the peak frequency above 2.5 Hz. We saw that the peak frequency increased significantly with *φ* beyond LFP-Power (2-way ANOVA with LFP-Power and *φ*, p<0.0001). The peak frequency below 2.5 Hz did not change significantly with *φ*. However, the peak frequency width appeared to narrow with *φ*, indicating that oscillations became more regular. We saw that the peak frequency width decreased significantly over all unanesthetized experiments both below 2.5 Hz (2-way ANOVA with LFP-Power and *φ*, p<0.0001) and above 2.5 Hz (2-way ANOVA with LFP-Power and *φ*, p<0.0001), indicating that both facets of the frequency spectra became more regular.

## Discussion

We applied ICA to recordings from rat auditory cortex and consistently found two components that alternate within synchronized states. The BL5 component has a high amplitude and follows the overall activity. The SUB component generates a theta oscillation from below the cortex. Cortical UP state activity varies depending on which component is active. These results indicate that there are two modes of operation within synchronized states.

### Related studies on differences within synchronized states and application of ICA

There are several other studies that note different modes within synchronized states. Recently, Miyawaki et al., (2017) describe low-amplitude synchronized states (LOW) seen in the hippocampus, which builds on previously noted small-irregular activity. LOW states may be related to our SUB-dominant states since UP state CSD amplitude is lower during SUB-dominant states. Notably, LOW states occur simultaneously in the hippocampus, entorhinal cortex, neocortex, and thalamus. Therefore, if there is a correlation between LOW states and SUB-dominant states, then low-amplitude irregular activity should be seen in the hippocampus during SUB-dominant states and SUB theta may not be easily noticed in the hippocampal LFP without using ICA due to contamination with other signals. Thus, our study hints that there is underlying theta oscillations during low-amplitude states in the hippocampus.

Although rat hippocampal theta oscillations have only been noted during desynchronized states in most studies, there are a few studies that look at theta oscillations during synchronized states. Hippocampal theta is seen in non-REM sleep during transition-to-REM states shortly before REM starts (Benington et al., 1994; Robert, Guilpin, & Limoge, 1999). However, transition-to-REM lasts up to 10s (Gottesmann, 1996), whereas SUB-dominant states can last significantly longer and do not necessarily lead to REM sleep. Several studies have noted that theta power in the hippocampus and thalamus during non-REM sleep can be as strong as theta power during REM sleep (Gaztelu, Romero-Vives, Abraira, & Garcia-Austt, 1994; Pedemonte, Gambini, & Velluti, 2005), but these oscillations may be more difficult to observe because of ongoing irregular activity. Finally, theta oscillations can be seen with increased tiredness during waking states (Vyazovskiy & Tobler, 2005).

ICA is a form of analysis that separates overlapping signals in a set of recordings (Amari, Cichocki, & Yang, 1996; Bell & Sejnowski, 1995; Brown et al., 2001; Herreras et al., 2015; A. A.Hyvärinen & Oja, 2000; A.Hyvärinen & Oja, 1997). Previously, ICA has been used a great deal in artifact removal as well as identifying regions of interest in fMRI and EEG (Calhoun & de Lacy, 2017; Eichele, Calhoun, & Debener, 2009; Esposito & Goebel, 2011; Huster, Plis, & Calhoun, 2015; McKeown, Hansen, & Sejnowsk, 2003; Onton, Westerfield, Townsend, & Makeig, 2006). More recently, ICA was used in LFP recordings to parse population involvement in the hippocampus. (Agarwal et al., 2014a; Benito, Martin-Vazquez, Makarova, Makarov, & Herreras, 2016; Herreras et al., 2015; Makarov et al., 2010). To our knowledge, this is the first study to use ICA to study synchronized states.

### Interpretation of results with ICA and limitations

One reason why ICA has not been used more on LFP recordings to date may be because LFP recordings have access to relatively detailed information, so there are more options available for analysis than fMRI and EEG. At the same time, ICA is limited in the fact that the number of components it can return is limited by the number of recordings. Because there are potentially more populations detected by neural recordings than can possibly be parsed, it has been unclear how ICA would behave when faced with real neural data. Recent simulations in Głąbska et al. (2014) and Makarov et al., (2010) show that ICA tends to group similar populations together, which is consistent with our results. With more recordings, ICA would be able to pick out more sources at finer resolution (Agarwal et al., 2014b; Jun et al., 2017).

As ICA groups similar populations with similar spatial profiles together, BL5 and SUB may not stand for single neural populations. In fact, ICA will parse any signal that affects recordings, including glia and movement artifacts. ICA components also may be composed of sets of synapses onto cells acting synchronously (Herreras et al., 2015). Furthermore, it is possible that two populations that act independently at some time points, and synchronously at others, may be separated into three components: one component for each of the populations individually and a third for the combination (Herreras et al., 2015).

Although we consistently found a SUB component which exhibited subcortical theta oscillations, the theta oscillations were more difficult to see in unanesthetized experiments compared to anesthetized experiments during synchronized states. The theta oscillation was often mixed with larger amplitude activity in unanesthetized experiments, while the theta peak was isolated in all anesthetized experiments. The theta peak may have been clearer for anesthetized SUB components vs. unanesthetized SUB components because: (1) activity under anesthesia is simpler than activity without anesthesia and so theta could have a dedicated component in anesthetized experiments, or (2) the SUB ICA component in unanesthetized experiments included movement artifact which allowed neocortical signal to leak into the component. In the first case, the SUB component may more closely reflect signals volume propagated from the hippocampus, which is directly below the auditory cortex and is known to show mixed frequencies during synchronized states (Buzsáki, 2002). On the other hand, in anesthetized experiments, signals volume propagated from below the cortex may have been split into multiple components if activity under anesthesia does not have as many independent signal generators as unanesthestized activity. Then it is possible that there are separate components in anesthetized experiments that account for subcortical theta activity as well as small- and large-irregular hippocampal activity.

Another limitation of our study is that instead of sleep staging, we compared ICA components to sleep stage parameters. Thresholds for these parameters are based on the data from rat recordings lasting 1-1.5 hours, which may not include all sleep stages. This means that sleep regions may not necessarily line up with sleep stages seen in other experiments. While our analysis does not provide a direct comparison with sleep stages, a comparison to the parameters allows us to see how BL5 and SUB signals relate to other states regardless of how sleep stages may be decided. Likewise, while there is no exact threshold for LFP-Power to determine synchronized vs. desynchronized states, we are still able to compare LFP-Power with BL5 and SUB signals.

### Possible implications for synchronized states and further research

Our results show that BL5 UP states have relatively high amplitude, while SUB UP states have lower amplitude. Amplitude during sleep has previously been characterized by slow-wave activity (SWA), which is high at sleep onset and slowly subsides throughout the night (Aeschbach & Borbély, 1993; Dijk, Brunner, & Borbély, 1990; Rodriguez et al., 2016; Vassalli & Dijk, 2009). Because of this change, we hypothesize that BL5-dominated states occur earlier during sleep, while SUB-states occur later, consistent with LOW states (Miyawaki et al., 2017). According to the two-process model of sleep, early sleep is associated with homeostatic processes along with SWA (Achermann, Dijk, Brunner, & Borbély, 1993; Borbély, 1982; Borbély, Daan, Wirz-Justice, & Deboer, 2016). At the same time, sleep is hypothesized to serve two functions: synaptic homeostasis and memory consolidation (Diekelmann & Born, 2010; Rasch & Born, 2013; Vassalli & Dijk, 2009). In particular, late sleep and lower amplitude stage 2 sleep has been shown to benefit certain types of memory consolidation (Diekelmann & Born, 2010; Hutchison & Rathore, 2015; Rasch & Born, 2013). Since BL5-dominated states occur with higher SWA, we propose that BL5-dominated states may play a role in homeostatic processes, while SUB states may be linked to memory consolidation.

If BL5 and SUB reflect different modes within synchronized states, then what populations could they represent? BL5 encompasses much of the cortex, is relatively high amplitude, and mimics the CSD. This means that BL5 could comprise a large population within the cortex, a high amplitude population, or both. Early work shows that glial recordings have relatively high amplitude and follow the LFP (Amzica & Steriade, 1998). Glia are also known to be involved in homeostatic processes during sleep (Bjorness et al., 2016; Clinton, Davis, Zielinski, Jewett, & Krueger, 2011; de Andrés, Garzón, & Reinoso-Suárez, 2011b; Fellin, Ellenbogen, De Pittà, Ben-Jacob, & Halassa, 2012; Poskanzer & Yuste, 2011). Furthermore, recent work with calcium imaging shows heavy glial involvement prior to neuronal involvement during SWA (Szabó et al., 2017). Therefore, BL5 may include glia.

SUB represents a population exhibiting subcortical theta oscillations. Theta oscillations are clock-like sinusoidal oscillations previously reported in the hippocampus along with other limbic structures (Buzsáki, 2002). Although the hippocampus lies directly below the rat auditory neocortex, we cannot be sure that SUB theta is hippocampal in origin since it is possible for LFP and therefore CSD signals to propagate. At the same time, the theta oscillations we observed had a lower frequency, matching type 2 theta oscillations previously associated with sensory processing and emotionally salient stimuli during waking states (Bland & Oddie, 2001; Tendler & Wagner, 2015). If type 2 theta oscillations are associated with emotionally relevant memory processing, then they could also be linked to hippocampal-amygdala co-activation (Hutchison & Rathore, 2015; Pelletier & Paré, 2004). It has also been hypothesized that theta oscillations correspond to information coming in to the hippocampus, while sharp waves indicate information transferred out of the hippocampus (Hasselmo, 1999). SUB theta oscillations during synchronized states may mean that there are low-amplitude theta oscillations within the hippocampus at the same time as sharp waves, where the theta oscillations stand for information coming in from another structure such as the amygdala, while information is begin transferred out via sharp waves.

Further research would involve verification of the roles of various populations in BL5 and SUB states. Being able to focus on one state within synchronized states may also help to shed light on not only mechanisms for oscillatory activity, but functional roles for activity. For instance, memory processing has been shown to be hampered from sleep deprivation for the latter half of the night (Rasch & Born, 2013). Previously, this was tied to a lack of REM sleep which plays a larger role later in the night. However, stage 2 non-REM sleep frequently alternates with REM in the latter half of the night and further study may elucidate a unique role for this stage in memory processing (Diekelmann & Born, 2010; Hutchison & Rathore, 2015; Rasch & Born, 2013).

We may also use this analysis to find different modes of operation, and specifically theta oscillations, in other species. For example, while rat hippocampal theta is easy to pick out, theta oscillations tend to be more difficult to identify and study in other species, such as humans and bats (Jacobs, 2013).

### Conclusion

ICA reveals that theta oscillations exist within synchronized states, and that they alternate with a high-amplitude population in the neocortex. These populations alternate on the same time scale and are associated with different neocortical activity. Therefore, these data indicate that synchronized states are not uniform, but may be separated according to at least two distinct functional roles.

## Acknowledgements

We would like to thank Tansi Khodai for performing the anesthetized experiments, and Austin Brockmeier for helpful discussions with ICA. Unanesthetized data was originally published in Sakata & Harris (2009) and Sakata & Harris (2012). This work was supported by the RIKEN Brain Science Institute (EM and TT), JSPS KAKENHI Grant Number JP18H05432 (TT), AMED under Grant Number JP18dm020700 (TT), Medical Research Council (MR/J004448/1 to SS), and Biotechnology and Biological Sciences Research Council (BB/K016830/1 and BB/M00905X/1 to SS).

## Author Contributions

Conceptualization: EMK. Methodology: EMK and TT. In vivo experiments: SS. Formal analysis: EMK. Resources: TT and SS. Writing - original draft: EMK. Writing - review & editing: EMK, TT, and SS. Supervision: TT.

---

**Supplementary Figure 1.**
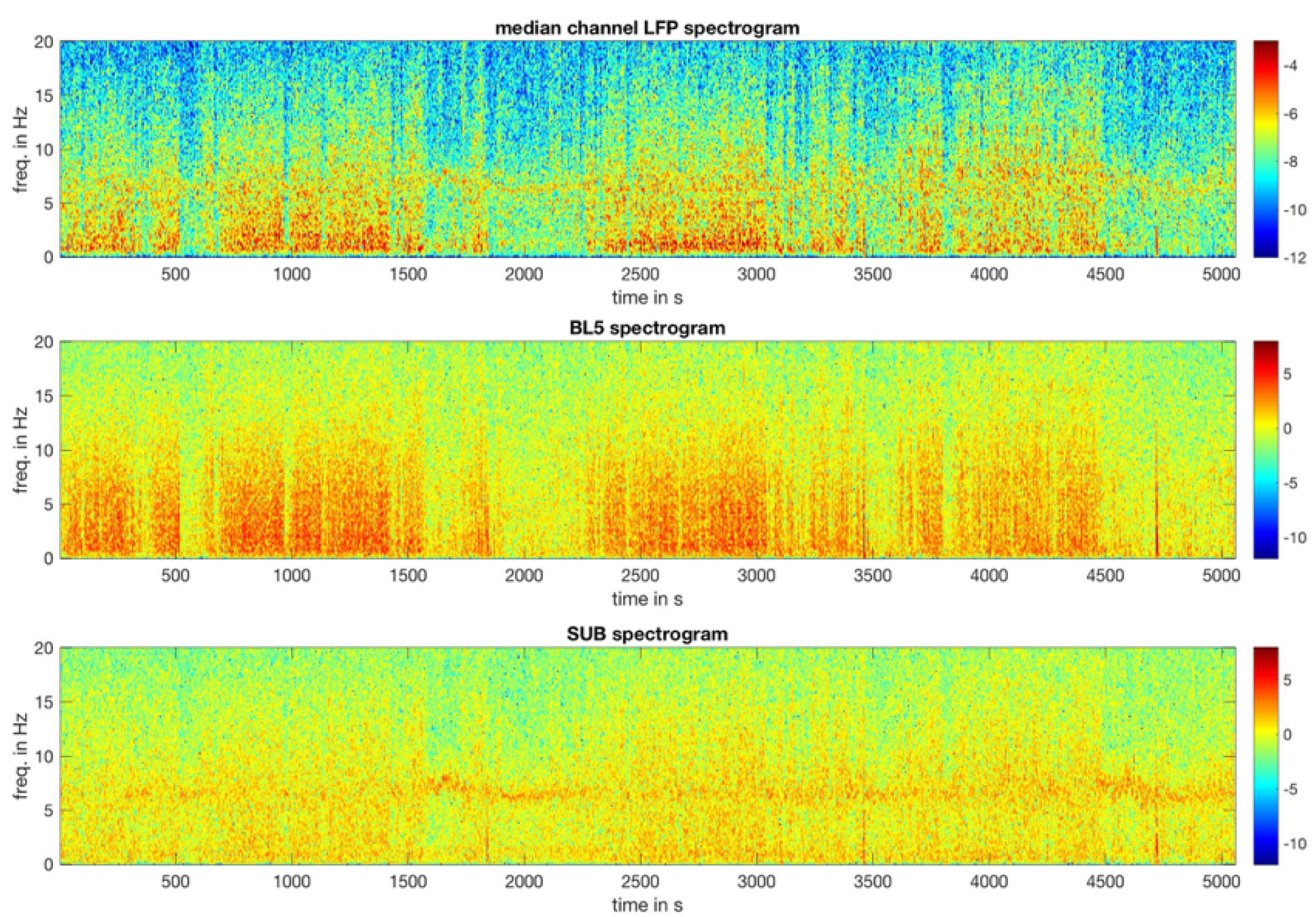
Spectrograms of an example unanesthetized experiment. The top panel shows the median spectrogram of the LFP recorded over all channels. The LFP-Power is the 1-5 Hz power of this spectrogram. The BL5 spectrogram shows strong low frequency oscillations when BL5 is active. The SUB spectrogram exhibits a theta oscillation which can sometimes be seen in the median LFP spectrogram. Notably, SUB exhibits a theta peak in the latter half of the experiment, which does not appear so clearly in the median LFP spectrogram. The SUB scaled amplitude shows that this peak is significant even when LFP-Power is high.

**Supplementary Figure 2.**
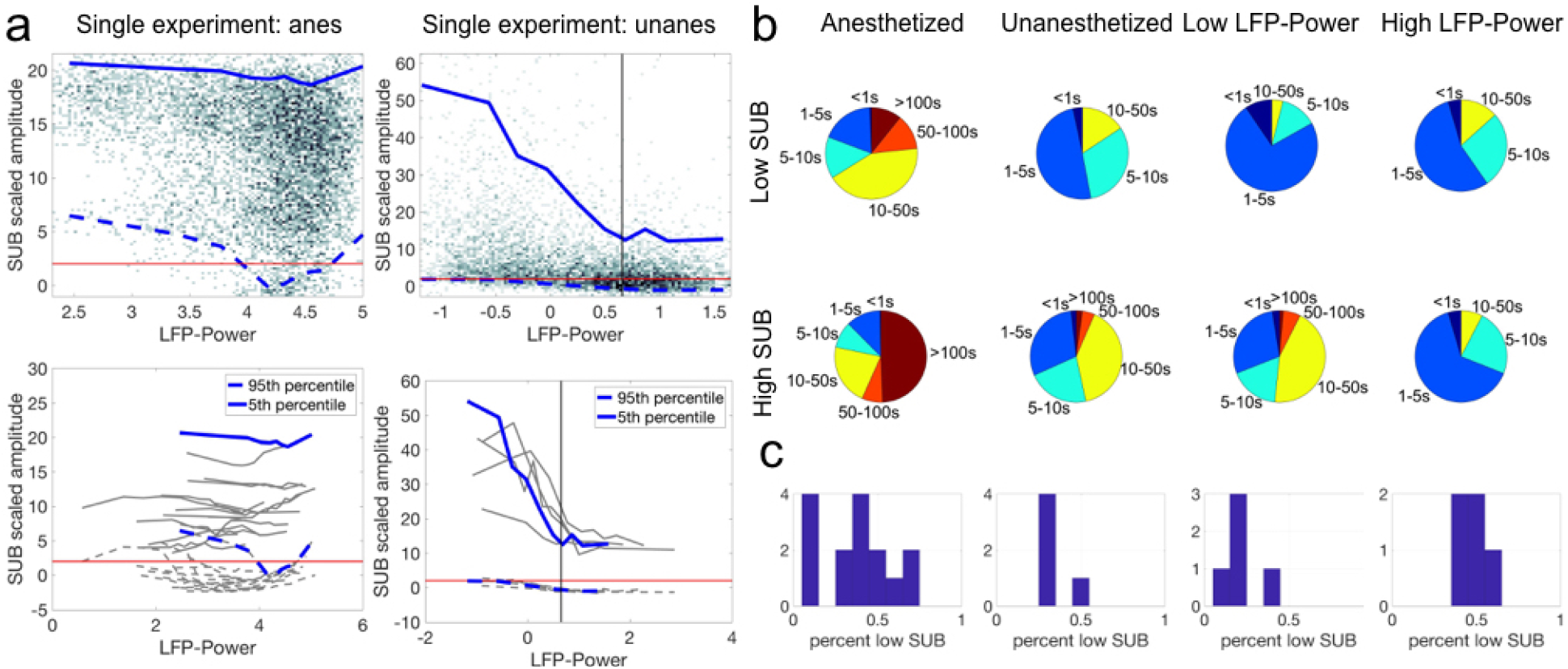
Relationship between SUB scaled amplitude and LFP-Power. (a) Similarly to φ, there is a wide range of SUB scaled amplitude over high values of LFP-Power. Top plots show density of LFP-Power vs. SUB scaled amplitude along with 5-95 percentile range outlined in blue. Bottom plots show the range calculated for all experiments, with example experiments highlighted in blue. Black vertical lines indicate threshold for low vs. high LFP-Power used to calculate state durations and time percentages. Red horizontal lines indicate threshold SUB scaled amplitude of 2. (b) State durations of low vs. high LFP-Power and low vs. high SUB scaled amplitude. (c) Percentage of time spent with low SUB scaled amplitude. Note that SUB scaled amplitude is significant approximately 50% of time during high LFP-Power.

**Supplementary Figure 3.**
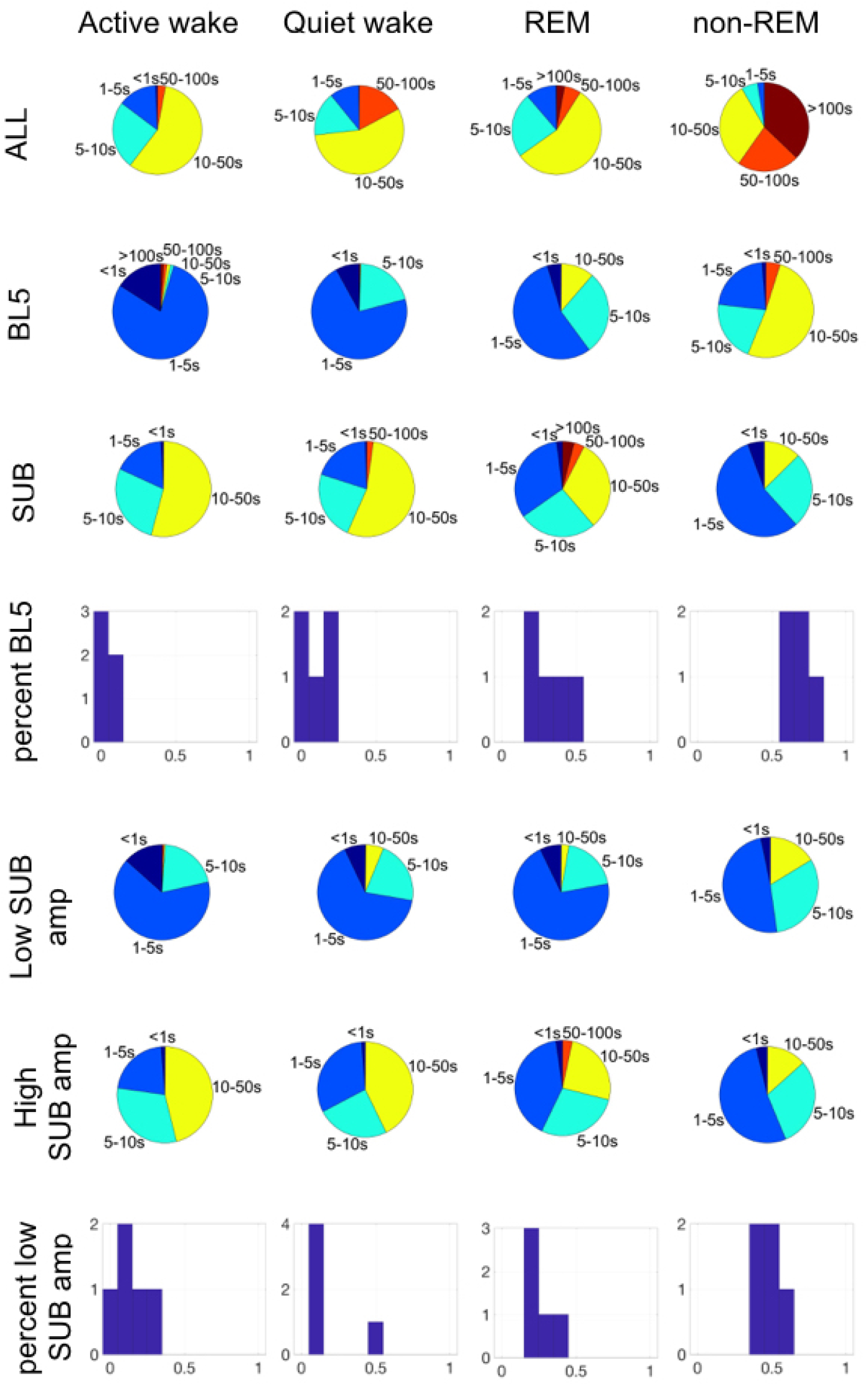
Time durations with respect to sleep parameters. Not that approximately 70% of time in non-REM regions is BL5-dominant, while 30% is SUB-dominant, similarly to states with high LFP-Power. Likewise, approximately 50% of time in non-REM regions has significant SUB scaled amplitude.

**Supplementary Table 1.** Statistical tests accompanying histograms in Fig. 1de. Matlab ttest() used throughout. DF=degress of freedom. STD=standard deviation.

| CONDITION | TEST | N | P-VALUE | DF | TEST<br>STATISTIC | MEAN | STD |
| --- | --- | --- | --- | --- | --- | --- | --- |
| CORRELTION BETWEEN LFP-POWER AND BL5<br>SCALED AMP, ANES | t-test | 17 | 7.4863E-07 | 16 | 7.8163 | 0.5478 | 0.2890 |
| CORRELTION BETWEEN LFP-POWER AND BL5<br>SCALED AMP, UNANES | t-test | 5 | 2.2632E-04 | 4 | 12.6679 | 0.7659 | 0.1352 |
| CORRELTION BETWEEN SMUA LFA AND BL5<br>SCALED AMP, ANES | t-test | 17 | 1.1480E-05 | 16 | 6.2555 | 0.4584 | 0.3022 |
| CORRELTION BETWEEN SMUA LFA AND BL5<br>SCALED AMP, UNANES | t-test | 5 | 0.0062 | 4 | 5.2803 | 0.4686 | 0.1984 |
| CORRELTION BETWEEN LFP RELATIVE THETA<br>AND SUB SCALED AMP, ANES | t-test | 15 | 0.0260 | 14 | 2.4884 | 0.1398 | 0.2175 |
| CORRELTION BETWEEN LFP RELATIVE THETA<br>AND SUB SCALED AMP, UNANES | t-test | 5 | 0.0532 | 4 | 2.7161 | 0.1298 | 0.1069 |
| CORRELTION BETWEEN SMUA RELATIVE THETA<br>AND SUB SCALED AMP, ANES | t-test | 15 | 0.4623 | 14 | 0.7558 | 0.0286 | 0.1465 |
| CORRELTION BETWEEN SMUA RELATIVE THETA<br>AND SUB SCALED AMP, UNANES | t-test | 5 | 0.5536 | 4 | 0.6459 | 0.0064 | 0.0223 |
| MEDIAN CORRELATION BETWEEN CSD L5 AND<br>BL5 WHEN ACTIVE, ANES | t-test | 17 | 3.5977E-12 | 16 | 18.3486 | 0.6233 | 0.1401 |
| MEDIAN CORRELATION BETWEEN CSD L5 AND<br>BL5 WHEN ACTIVE, UNANES | t-test | 5 | 1.3056E-04 | 4 | 14.5274 | 0.6595 | 0.1015 |
| MEDIAN CORRELATION BETWEEN SMUA AND<br>BL5 WHEN ACTIVE, ANES | t-test | 17 | 1.8073E-10 | 16 | -14.1601 | -.3716 | 0.1082 |
| MEDIAN CORRELATION BETWEEN SMUA AND<br>BL5 WHEN ACTIVE, UNANES | t-test | 5 | 0.0027 | 4 | -6.5997 | -.2018 | 0.0684 |
| MEDIAN CORRELATION BETWEEN CSD L5 AND<br>SUB WHEN ACTIVE, ANES | t-test | 15 | 9.7414E-05 | 14 | 5.3779 | 0.1468 | 0.1057 |
| MEDIAN CORRELATION BETWEEN CSD L5 AND<br>SUB WHEN ACTIVE, UNANES | t-test | 5 | 1.3252E-04 | 4 | 14.4726 | 0.4246 | 0.0656 |
| MEDIAN CORRELATION BETWEEN SMUA AND<br>SUB WHEN ACTIVE, ANES | t-test | 15 | 0.4033 | 14 | -.8619 | -.0154 | 0.0690 |
| MEDIAN CORRELATION BETWEEN SMUA AND<br>SUB WHEN ACTIVE, UNANES | t-test | 5 | 0.1243 | 4 | -1.9408 | -.0353 | 0.0406 |

**Supplementary Table 2.**
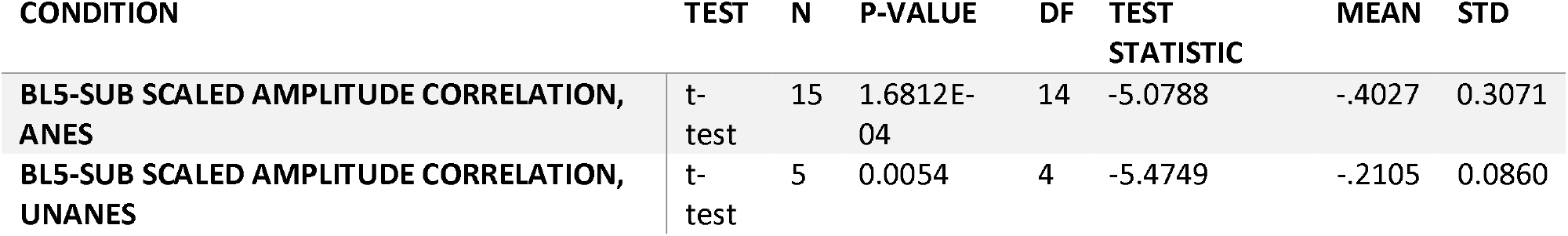
Statistical tests accompanying histograms in Fig. 2aii. Matlab ttest() used throughout. DF=degrees of freedom. STD=standard deviation.

| CONDITION | TEST | N | P-VALUE | DF | TEST<br>STATISTIC | MEAN | STD |
| --- | --- | --- | --- | --- | --- | --- | --- |
| BL5-SUB SCALED AMPLITUDE CORRELATION,<br>ANES | t-test | 15 | 1.6812E-04 | 14 | -5.0788 | -.4027 | 0.3071 |
| BL5-SUB SCALED AMPLITUDE CORRELATION,<br>UNANES | t-test | 5 | 0.0054 | 4 | -5.4749 | -.2105 | 0.0860 |

**Supplementary Table 3.**
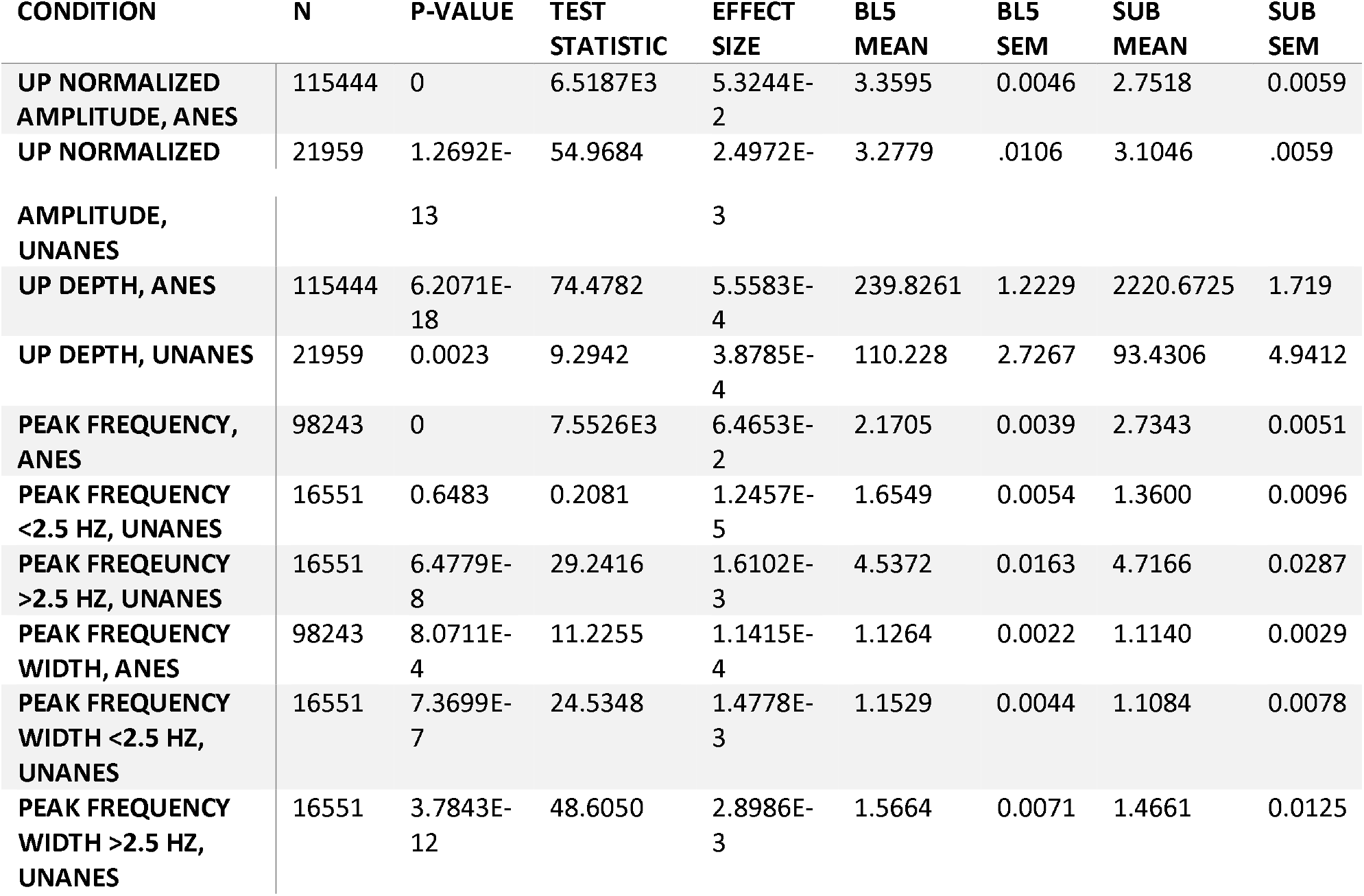
Statistical tests accompanying results in Fig. 5. Matlab anovan() used throughout. Means and standard error of the mean (SEM) calculated with multcompare(). Effect size is the sum of squares divided by total sum of squares.

